# Rational Decision On the Use of Antibiotics During the Dry Period in Dairy Cows

**DOI:** 10.1101/667873

**Authors:** Luis O. Lopes, Anna M. C. Lima

**Affiliations:** Doctor in Veterinary Sciences at the Federal University of Uberlândia, state of Minas Gerais, Brazil; Professor at the Faculty of Veterinary Medicine, Federal University of Uberlândia - FAMEV-UFU, Uberlândia, state of Minas Gerais, Brazil

## Abstract

The aim of this study was to evaluate the use of antibiotics in cows during the dry period. The survey was performed on 148 teats during the dry period, with sample collection in the period D-70 (70 days before delivery) and D14 (14 days after delivery). The milk samples were collected for the Strip Cup Test (SCT), California Mastitis Test (CMT), Microbiological Culture, Somatic Cell Count (SCC), Somatic Cell Score (SCS) and Hyperkeratosis (HK). The groups in which there were no microorganisms grow were divided into two groups, in the first group only the internal sealant in the teat was used (Group 1) and there was another group with the intramammary antibiotic use associated with the internal sealant (Group 2). Teats which were considered positive, with microbiological growth, were treated with the intramammary antibiotic associated with the internal sealant (Group 3). In the comparison of the results of the CMT test between D-70 and D14, a statistical difference was observed in Groups 2 and 3. Group 3, which comprises the positive teats in D-70 presented a reduction of 83.87% and 32.26% in the CMT test between D-70 and D14. Regarding HK, group 1 and 2 had a statistical difference in relation to group 3 in D-70 and D14. As for the numbers of bacteria isolated in D-70 and D14, there was no difference comparing Group 1 and Group 2, unlike Group 3, which had a difference. Group 1 and Group 2 were all negative teats in D-70, showing that the intramammary antibiotic did not influence the outcome of D14. In group 3 there was a reduction of isolates from 62 to 15 in D14. The most prevalent microorganism was Streptococcus agalactiae with 43.37% of the total isolates, followed by Staphylococcus aureus (16.87%) and Corynebacterium spp. (13.25%) and Coagulase negative Staphylococcus (SCN) (10.84%). The selective treatment of teats in dry dairy cows has advantages over Blanket Dry Cow Therapy by reducing the indiscriminate use of antibiotics, avoiding bacterial resistance, ensuring better milk quality and greater food safety. Antibiotics should only be used for teats with subclinical mastitis, with the microbiological culture at the end of lactation performed by fourth individual mammary.

## Introduction

Mastitis is the most frequent disease which affects dairy herds causing prejudice worldwide ^(1)^, either due to reduced milk production, total or partial loss of the mammary gland, risk of transmission to other animals or even the animal’s death ^(2)^. The cause of mastitis is multifactorial, it can be influenced by the environment in which the animal is found, the affected microorganism and the response of each individual to the causative agent ^(3–5)^.

There is a trend towards the rational use of antibiotics, both for reducing residues in animal products for food safety and also for decrease of the microorganisms’ resistance to antibiotics used in animals ^(6)^. The reduction in the use of antibiotics is important for the efficiency of treatments in cases of diseases ^(7)^.

In dairy cows, besides the use of antibiotics in the treatment of clinical and subclinical mastitis during lactation, the use of antibiotics for intramammary use at the time of drying among lactations is also used, but without the proper diagnosis of mastitis, with application of antibiotics in all teats and generally with association with internal sealant, acting as mechanical barrier against environmental agents in the pre-labour ^(8,9)^.

Mastitis is a complex and widespread disease with multiple microorganisms’ etiology and conditioned by several factors, added to the specificity of treatment to each of the agents, it becomes even more difficult to treat, requiring studies that evaluate the best treatment in the period of drying, to perform the selective treatment in dairy cows (STDC) ^(10,11)^.

Legislation on the use of antibiotics in disease prevention is increasingly restricted in the area of animal production, including the treatment of antibiotics in the drying period ^(12,13)^. There is a concern of decreasing the use of antibiotics in the drying of dairy cows, treating only the teats with intramammary infections with the previous diagnosis.

During the drying period in dairy cows, there is no technical standard in decision making, some professionals choose to treat all teats with antibiotics regardless of historical and mammary gland health assessment (BDCT), other professionals and producers use internal sealants or antibiotics in teats according to the record of SCC and clinical mastitis of the animal, justifying studies to clarify the best management of cows at the end of lactation at each farm.

Many studies on the selective treatment in the drying period consider the cow with a history of SCC during lactation ^(8,9,14)^. It can be noticed that there is a need for further studies with individual evaluation of mammary quarters for an even greater reduction in the use of antibiotics ^(15)^.

The objective of this research was to evaluate the use of antibiotics in cows during the drying period.

## Materials and methods

### Herd characterization

The study was carried out on a dairy farm, with a latitude of 17° 50 ′07 ″S and a longitude of 46° 30’ 52” W, with a daily average production of 9,000 liters of milk, two milking per day, with approximately 450 dairy cows, the average per cow of 30 kg per day in 3 milking, with blood level ranging from 1/2 to 31/32 Dutch/Gir, with semi-intensive breeding system, throughout the year. All animals within the same farm were submitted to the same management, with silage-based feed and concentrate throughout the year, varying proportions according to their milk production, lactation days and days in gestation (NRC, 2001).

The primiparous cows, the ones with antibiotic application or case of clinical mastitis were discarded in the last 30 days before drying, presence of lumps on the D-70 Strip Cup Test (70 days before delivery), presence of conditions in the hull, drying period less than 50 days and greater than 90 days before delivery.

### Trial design

The research was performed in 37 cows (148 teats) in the drying period, for six months, with sample collection on D70 period (70 days before delivery) and D14 (14 days after delivery), with sampling by teat.

Drying of the dairy cows was performed on average 60 ± 7 days before the expected date of delivery, with the type of treatment in each group according to the results of the tests in D-70, 10 days before drying.

The milk samples were collected for the Strip Cup Test (SCT), CMT (California Mastitis Test), microbiological culture, SCC (somatic cell count), SCS (Somatic Cell Score), hyperkeratosis (HK) in each animal on D-70 and D14, according to Figure 1.

**Figure 1.**
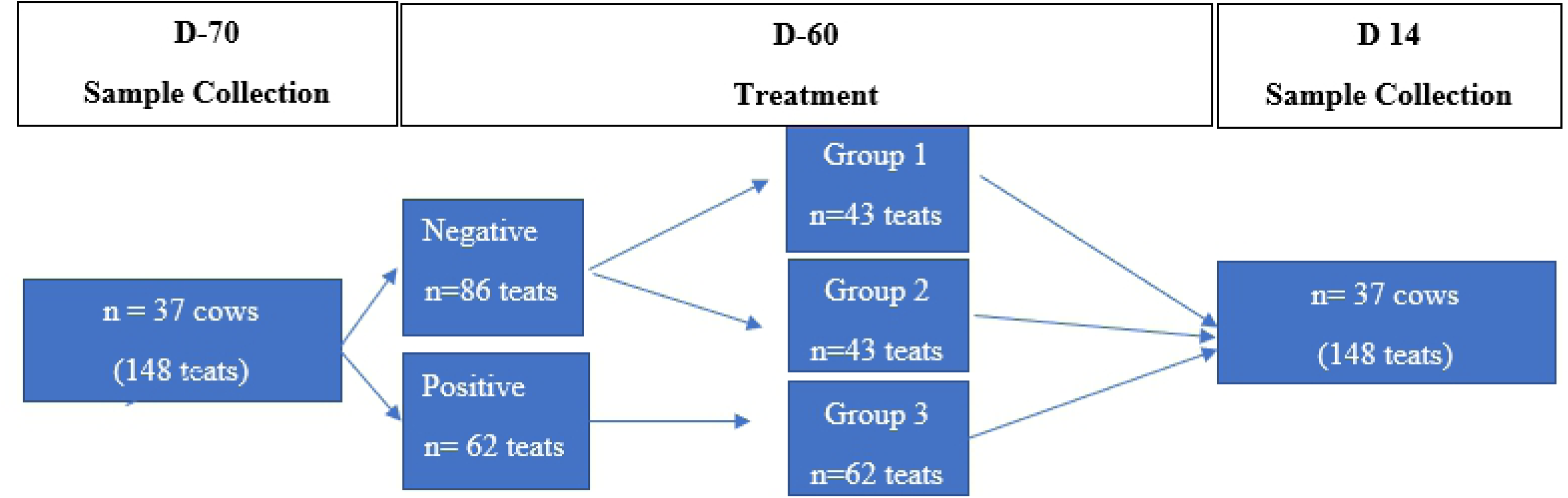
Description of procedures performed before (D-70 D-60) and after delivery(D14).

Negative teats were considered as those whose milk samples did not show growth of microorganisms, and the positive teats were the milk samples that showed growth of microorganisms from the microbiological culture test performed on D-70.

The 148 teats were divided into three groups after the results of the examinations on D-70. The teats considered negative had two different treatments, in the first one only the internal sealant was used in the teat (Group 1), and another group with the intramammary antibiotic associated with the internal sealant (Group 2). The teats considered positive were treated with the intramammary associated with the internal sealant (Group 3).

The intramammary antibiotic used was based on Cloxacillin Benzathine 600 mg, using one tube per teat (3.6 g), in groups 2 and 3. In all teats, in the 3 treatment groups, one tube was used per internal sealant teat based on bismuth subnitrate (4 g) to reduce the influence of the environment on the health of the mammary gland during the dry period.

The use of the sealant in all the teats was to avoid external contamination of the teats during the dry period, to better evaluate the dynamics of the results in the established groups.

In the application of the products at the time of drying (D-60), the teats were disinfected with 10% topical polyvidone iodine solution and paper towel drying after 30 seconds, performing another antisepsis of the teat sphincter using alcohol-wetted cotton to 70%. After the disinfection of the teats, the cannula of the antibiotic tubes was introduced, depending on the treatment group, massaging the teat to propel the substance upwards and soon after the internal sealant tube was applied via cannula inside the canal of the teat.

### CMT Test

For the CMT, 2 mL of milk from each mammary quarter were removed on the CMT plate, to which was added, in the same proportion as the CMT solution, an anionic detergent (alkyl lauryl sodium sulfate), homogenized with circular movements for 10 seconds, prior to reading, which is determined by the sample gelation, if the test is positive.

The interpretation of the test is usually classified as negative, one cross, two crosses or three crosses, according to absence or increase of the viscosity from the mixture (17,18). The mammary quarters that presented viscosity from the score of a cross were considered positive on this research.

The animals were evaluated by performing the CMT Test, on D-70 and D14, always performed by the same person to avoid subjectivity to the test.

### Hyperkeratosis score in the teats (HK)

The severity of hyperkeratosis, an excessive growth of the teat skin, was classified visually through an evaluation in scores, ranging from 0 (normal) to 3 ^(19)^. This tool has as the main objective the aid of problem identification, which among the main causes is the proper functioning of the milking equipment and correct management of milking in dairy herds.

### Somatic Cell Count

In order to perform the Somatic Cell Count (SCC), milk samples were collected in plastic bottles containing bronopol preservative on D-70 and D14 and sent on the same day of collection to the laboratory.

Bronopol is a tablet-form preservative which in contact with milk results in a light pink-colored blend. For the SCC determination of the milk samples, the flow cytometry technique was applied using the SOMACOUNT 500 device, with a result expressed the number of cells multiplied by 10^3^ / mL of milk (20). In the SCC results, it is usually considered an animal with subclinical mastitis presenting values > 200,000 cells / mL of milk (ECS>4) (21).

In order to linearize the data, SCC was transformed into Somatic Cell Score (SCS), with SCS = [log2 (SCC / 100)] +3 (22,23).

### Microbiological Culture

On D-70 and D14, after the Strip Cup and CMT tests, milk samples collections were carried out for individual microbiological diagnosis in all teats, aiming at the identification of pathogens causing mastitis. The milk samples were obtained immediately prior to milking, after discarding the first three milk jets.

It was disinfection of the teats with 10% topical polyvidone iodine solution and drying with paper towels after 30 seconds was performed. At the time of collection, the antisepsis of teat sphincter was held using cotton wool moistened with 70% alcohol. The milk samples were collected from individual teats and placed in sterile bottles previously identified with the number and teat of the animal. The sampled material was frozen and then sent in an isothermal container with recyclable ice to the laboratory for microbiological diagnosis with the isolation and characterization of the microorganisms.

Samples of milk were cultured in 5% (v / v) blood agar plates, incubated in aerobiosis in a bacteriological heating chamber at 37°C and analyzed after 24 and 48 hours. Following the incubation, the growth characteristics of the colonies on blood agar were recorded, such as hemolysis production, pigment, type of development and pigmentation of the colonies, observing hereinafter the morpho-tinctorial characters using the Gram staining technique.

Colonies which demonstrated to be Gram-positive cocci were submitted to catalase and slow coagulase tests with rabbit plasma. The readings for the verification of coagulase production were performed one, two, three, four and 24 hours after incubation of the samples at 37 ° C ^(24)^. Biochemical tests were performed to identify isolated microorganisms ^(25)^.

The catalase and coagulase positive strains were tested for acetoin production with the Methyl Red and Voges-Proskauer test (MRVP broth) for the differentiation of Staphylococcus aureus and another coagulase-positive Staphylococcus. The acetoin-producing strains were tested for whether or not maltose and trehalose could be used ^(26)^. Strains that presented positive results for these tests were classified as *Staphylococcus aureus* ^(27,28)^

### Statistical analysis

For the comparison among the three groups with HK, the Tukey’s test was used. By comparing the results of D-70 and D14 analyses with the CMT test, the Chi-Square and T-Test were performed on the HK results.

As for the numbers of isolates, the Chi-Square test was used to compare the results of the D-70 and D14 analyses among each group.

In all statistical tests, a significance level of 5% was applied, using ACTION 3.0 software ^(29)^.

### Economic analysis

For the analysis of the cost regarding the drying treatment of the cows, the real value of the examinations from this study was used, adding to the values of the medicines medicine. The values of microbiological culture tests were $2.05 per sample, SCC test $ 0.70 per animal, CMT test $0.03 per teat. The cost of the antibiotic was $3.07 and the sealant $2.25 per teat.

## Results and discussion

Comparing the results of the CMT test between D-70 and D14, a statistical difference was observed in Groups 2 and 3, according to Table 1. Group 3, which are positive teats on D-70, had a reduction of 83.87% and 32.26% in the CMT test between D-70 and D14, with statistical difference.

**Table 1.**
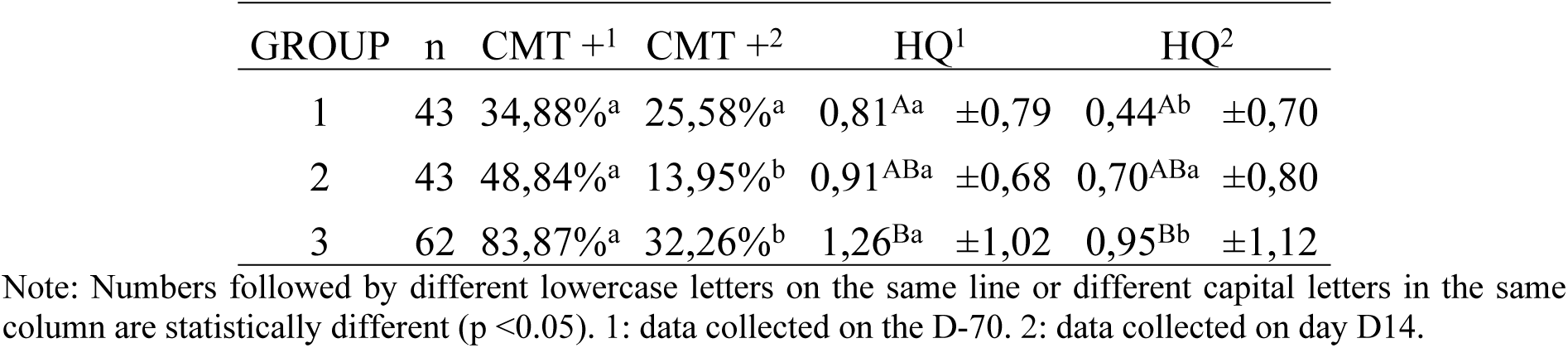
Mean and Standard Deviation of the CMT and HK test in each treatment group on D-70 and D14.

| GROUP | n | CMT +<sup>1</sup> | CMT +<sup>2</sup> | HQ<sup>1</sup> | HQ<sup>2</sup> |
| --- | --- | --- | --- | --- | --- |
| 1 | 43 | 34,88%<sup>a</sup> | 25,58%<sup>a</sup> | 0,81<sup>Aa</sup> ±0,79 | 0,44<sup>Ab</sup> ±0,70 |
| 2 | 43 | 48,84%<sup>a</sup> | 13,95%<sup>b</sup> | 0,91<sup>ABa</sup> ±0,68 | 0,70<sup>ABa</sup> ±0,80 |
| 3 | 62 | 83,87%<sup>a</sup> | 32,26%<sup>b</sup> | 1,26<sup>Ba</sup> ±1,02 | 0,95<sup>Bb</sup> ±1,12 |

Note: Numbers followed by different lowercase letters on the same line or different capital letters in the same column are statistically different (p <0.05). 1: data collected on the D-70. 2: data collected on day D14.

Intramammary antibiotic treatment during the dry period is recommended for treatment of subclinical mastitis, in association with internal sealant, as used in this research (11). The bismuth subnitrate used in this study is the only sealant base found in the market, with proven results in other studies (11,30).

The positivity of the CMT test in Groups 1 and 2 on D-70 in teats that did not show bacterial growth demonstrates the need for caution when using the test at the time of drying. The presence of microorganisms in negative teats may occur in the CMT Test, with low specificity (31,32).

The mean SCS values of the three treatment groups, before and after the dry period, were above 4 (greater than 200,000 cels / mL), even in groups 1 and 2 that did not have microbiological growth. For the selective treatment in drying, research has reported that SCS and CMT, may have errors in interpretations and interfere with the segregation of treatment in the dry period (31,33-36).

In the individual SCC, values below 200.000 cels / mL of milk refer to the animals without intramammary infection, but research shows that intramammary infection can occur even in values within the reference standards (37-39) and that the CMT Test, even with a negative result, may present microorganisms in the mammary quarter (31).

In Table 1, regarding HK, group 1 and 2 obtained a statistical difference in relation to group 3 on D-70 and D14. There was a difference in group 1 and 3 and no difference in group 2 comparing the results of each group on D-70 with D14, which shows that the dry period can decrease the degree of HK, but in some cases are irreversible, according to the injury level in the teat (19). The higher the HK score, the lower the mechanical barrier and the higher the ease of teat contamination, consequent incidence of subclinical mastitis ^(19,40,41)^.

As for the numbers of isolated bacteria on D-70 and D14, there was no difference comparing Group 1 and Group 2, but with a difference in group 3. Groups 1 and 2 were all negative teats on D-70, showing that the intramammary antibiotic did not influence the result of D14, according to Table 2. In group 3 there was a reduction of isolates from 62 to 15 on D14. Research has reported a reduction of 70.27% in infection in treated teats, comparing before and after delivery ^(15)^.

**Table 2.** The number of microorganisms isolated in milk samples in relation to the treatment groups on D-70 and D14.

|  | Groups |  |  |  |  |  | Total |  |
| --- | --- | --- | --- | --- | --- | --- | --- | --- |
|  | 1 |  | 2 |  | 3 |  |  |  |
|  | D-70 | D14 | D-70 | D14 | D-70 | D14 | n | % |
| *Corynebacterium sp.* | 0 | 1 | 0 | 0 | 10 | 0 | 11 | 13,25 |
| *Escherichia coli* | 0 | 0 | 0 | 0 | 2 | 2 | 4 | 4,82 |
| *Klebsiella pneumoniae* | 0 | 0 | 0 | 0 | 0 | 1 | 1 | 1,20 |
| *Staphylococcus aureus* | 0 | 0 | 0 | 1 | 5 | 8 | 14 | 16,87 |
| *Staphylococcus coagulase negative* | 0 | 2 | 0 | 2 | 5 | 0 | 9 | 10,84 |
| *Streptococcus agalactiae* | 0 | 0 | 0 | 0 | 32 | 4 | 36 | 43,37 |
| *Streptococcus bovis* | 0 | 0 | 0 | 0 | 3 | 0 | 3 | 3,61 |
| *Streptococcus uberis* | 0 | 0 | 0 | 0 | 5 | 0 | 5 | 6,02 |
| Total: | 0<sup>a</sup> | 3<sup>a</sup> | 0<sup>a</sup> | 3<sup>a</sup> | 62<sup>a</sup> | 15<sup>b</sup> | 83 | 100 |

Note: Numbers followed by different lowercase letters on the same line are statistically different ( $p < 0.05$ ).

The most prevalent microorganism, according to Table 2, on D-70 and D14 was *Streptococcus agalactiae* with 43.37% from the total isolates, followed by *Staphylococcus aureus* (16.87%) and *Corynebacterium spp.* (13.25%) and *Staphylococcus coagulase negative (SCN)* (10.84%), with a significant difference in the number of isolates, found on D-70 and D14. Surveys obtained a frequency of *Streptococcus spp*. (24.52%), *Staphylococcus aureus* (13.69%), *Corynebacterium spp.* (6.87%) and SCN (75.97%) ^(15)^. *Streptococcus spp.* (2.79%) was less frequent, there was no growth of *Staphylococcus aureus*. On the other hand, CNS (54.31%) were more frequent, followed by *Corynebacterium spp*. (12.69%), in the results of microbiological culture examinations per mammary quarter before and after delivery ^(42)^.

The bacteria *Corynebacterium spp., Staphylococcus coagulase negative, Streptococcus bovis* and *Streptococcus uberis* were bacteriologically cured in the treatment of group 3 between D-70 and D14, according to table 2. The frequency of *Streptococcus agalactiae* decreased from 32 to 4 (87.5%), and *Staphylococcus aureus* increased from 5 to 8 (60%) in the treatment of Group 3 on D-70 compared to D14. The result in the treatment of subclinical mastitis in positive teats in the dry period is directly related to the type of agent most prevalent in the herd ^(43)^.

Groups 1 and 2, comparing the number of isolated microorganisms on D-70 and D14, were not statistically different. The difference in the treatment in Groups 1 and 2 was the use of the antibiotic, reinforcing the possibility of the STDC in teats considered negative. In another study, they compared the same treatment with and without antibiotic, associated with the sealant, but in low-risk cows with a history of SCC below 200 x 103 cells / mL, and obtained the same result of negative teats with and without antibiotics, with prior segregation by SCC analysis history ^(44)^.

The economic viability of STDC is related to the incidence of mastitis in each herd, that is, the decrease in the use of antibiotics is directly related to the incidence of subclinical mastitis at the end of lactation within each herd ^(9)^.

Table 3 shows the estimated cost of treatment with the microbiological culture test, comparing with the hypotheses of segregation using the CMT, SCC and BDCT (Blanket Dry Cow Therapy) tests which use antibiotic in all teats regardless of the previous examination.

**Table 3.** The estimated cost of the selective treatment in the drying of dairy cows according to each diagnostic method in positive and negative teats.

| Método | n | Positivos |  | n | Negativos |  |  |
| --- | --- | --- | --- | --- | --- | --- | --- |
| | | CP (\$) | CTP (\$) | | CN (\$) | CTN (\$) | CF (\$) |
| Cultura | 83 | 7,37 | 611,71 | 65 | 4,30 | 279,50 | 891,21 |
| CMT | 88 | 5,35 | 470,80 | 60 | 2,28 | 136,80 | 607,60 |
| BDCT | 148 | 5,32 | 787,36 | 0 | 5,32 | - | 787,36 |

Note: CP: Cost per positive teat (exams + antibiotic + sealant), TCP: total cost in positive teats, CN: Cost per negative teat (exams + sealant), TCN: total cost in negative teats and FC: Final cost (positive and negative teats).

Treatment at the end of lactation using SCC history during lactation has been the most used in other studies (9,14,21,45), but microbiological growth may occur in cows with values below 200 × 10^3^ cells / mL (45). With the use of the microbiological culture as a parameter, the FC was $891.21, with the SCC test of $792.72 and with the use of the BDCT method was $787.36. The cost using culture test is variable, which depends on the contamination degree of the mammary gland, the antibiotic use varies, with the possibility of being more financially viable than the BDCT.

The estimated cost of treatment in positive teats, using the previous culture examination (CP), was $7.37, test of CMT $5.35 and BDCT $5.32. In negative teats (CN), the estimated cost of the culture test was $4.30, CMT test $2.28 and BDCT $5.32. When the CMT test was used to segregate positive and negative teats, the final cost (FC) was lower at the time of drying with $607.60, but there is a possibility of microbiological growth in teats considered negative (31,46), with fake positive occurrences in the test, in negative teats, as it occurred in this study.

Using the culture test at the end of the D-70 lactation, it has a greater number of negative teats diagnosed, using fewer antibiotics and reducing the cost of treatment in the dry period. Selective treatment at the time of drying using the culture method may be economically more viable than BDCT if the number of intramammary infections is low with lower intramammary antibiotic expense ^(9, 42)^.

There are several guidelines from international organizations recommending caution in the use of antibiotics in production animals in order to reduce residues in food and bacterial resistance in animals and also humans ^(7,13,47–50)^.

Both economically and in relation to the sanity of the fourth mammary during the dry period, the selective treatment of teats with the microbiological culture is recommended. The SCC test should not be used in the STDC at the end of lactation as the test results judge the animal rather than the individual teat, limiting the result in the antibiotic reduction protocol established in the drying of cows. It is important to provide a safety diagnosis on drying using the STDC to ensure a reduction in the use of antibiotics and sanity of the mammary gland during the dry period.

## Conclusion

The selective treatment of teats in the drying of dairy cows offer advantages over Blanket Dry Cow Therapy, reducing the indiscriminate use of antibiotics, avoiding bacterial resistance, as well as ensuring better milk quality and greater food safety. Antibiotics should only be used for teats with subclinical mastitis, with the microbiological culture at the end of lactation performed by individual mammary quarter.

## References

1. Ruegg PL. A 100-Year Review: Mastitis detection, management, and prevention. J Dairy Sci [Internet].Elsevier; 2017 Dec 1 [cited 2017 Dec 20];100(12):10381–97. Available from: http://linkinghub.elsevier.com/retrieve/pii/S0022030217310329. https://doi.org/10.3168/jds.2017-13023.

2. Akers RM. A 100-Year Review: Mammary development and lactation. J Dairy Sci [Internet]. 2017 Dec [cited 2017 Dec 20];100(12):10332–52. Available from: http://linkinghub.elsevier.com/retrieve/pii/S0022030217310354. https://doi.org/10.3168/jds.2017-12983.

3. Down PM, Bradley AJ, Breen JE, Green MJ. Factors affecting the cost-effectiveness of on-farm culture prior to the treatment of clinical mastitis in dairy cows. Prev Vet Med [Internet]. Elsevier; 2017 Sep 15 [cited 2017 Dec 20];145:91–9. Available from: http://linkinghub.elsevier.com/retrieve/pii/S0167587717302684. https://doi.org/10.1016/j.prevetmed.2017.07.00

4. Down PM, Green MJ, Hudson CD. Rate of transmission: A major determinant of the cost of clinical mastitis. J Dairy Sci [Internet]. Elsevier; 2013;96(10):6301–14. Available from: http://linkinghub.elsevier.com/retrieve/pii/S0022030213005559. https://doi.org/10.3168/jds.2012-6470

5. Shahid MQ, Reneau JK, Chester-Jones H, Chebel RC, Endres MI. Cow- and herd-level risk factors for on-farm mortality in Midwest US dairy herds. J Dairy Sci [Internet]. Elsevier; 2015;98(7):4401–13. Available from: http://linkinghub.elsevier.com/retrieve/pii/S0022030215003215. https://doi.org/10.3168/jds.2012-6470

6. Bressmann T. Self-inflicted cosmetic tongue split: a case report. J Can Dent Assoc [Internet]. 2004 Mar [cited 2018 Jan 1];70(3):156–7. Available from: https://amr-review.org/sites/default/files/160525_Final paper_with cover.pdf. https://doi.org/.1016/j.jpha.2015.11.005

7. Wells V, Piddock LJ V. Addressing antimicrobial resistance in the UK and Europe. Lancet Infect Dis [Internet]. Elsevier; 2017 Dec 1 [cited 2018 Jan 1];17(12):1230–1. Available from: http://www.ncbi.nlm.nih.gov/pubmed/29133171. https://doi.org/10.1016/j.jpha.2015.11.005

8. Higgins HM, Golding SE, Mouncey J, Nanjiani I, Cook AJC. Understanding veterinarians’ prescribing decisions on antibiotic dry cow therapy. J Dairy Sci [Internet]. American Dairy Science Association; 2017 Apr 1 [cited 2017 Dec 18];100(4):2909–16. Available from: http://linkinghub.elsevier.com/retrieve/pii/S0022030217300747. https://doi.org/10.3168/jds.2016-11923

9. Scherpenzeel CGM, Hogeveen H, Maas L, Lam TJGM. Economic optimization of selective dry cow treatment. J Dairy Sci [Internet]. American Dairy Science Association; 2017 Feb [cited 2017 Dec 18];101(2):1530–9. Available from: http://linkinghub.elsevier.com/retrieve/pii/S0022030217311098. https://doi.org/10.3168/jds.2017-13076

10. Dolder C, van den Borne BHP, Traversari J, Thomann A, Perreten V, Bodmer M. Quarter- and cow-level risk factors for intramammary infection with coagulase-negative staphylococci species in Swiss dairy cows. J Dairy Sci [Internet]. 2017 Jul [cited 2017 May 7];100(7):5653–63. Available from: http://linkinghub.elsevier.com/retrieve/pii/S0022030217303703. https://doi.org/10.3168/jds.2016-11639

11. Golder HM, Hodge A, Lean IJ. Effects of antibiotic dry-cow therapy and internal teat sealant on milk somatic cell counts and clinical and subclinical mastitis in early lactation. J Dairy Sci [Internet]. American Dairy Science Association; 2016 Sep 1 [cited 2017 Mar 1];99(9):7370–80. Available from: http://linkinghub.elsevier.com/retrieve/pii/S0022030216303745. https://doi.org/10.3168/jds.2016-11639

12. Kuipers A, Koops WJ, Wemmenhove H. Antibiotic use in dairy herds in the Netherlands from 2005 to 2012. J Dairy Sci [Internet]. 2016;99(2):1632–48. Available from: http://linkinghub.elsevier.com/retrieve/pii/S0022030215009054. https://doi.org/10.3168/jds.2014-8428

13. Piddock LJV. Reflecting on the final report of the O’Neill Review on Antimicrobial Resistance. The Lancet Infectious Diseases. 2016. p. 767–8. https://doi.org/10.1016/S1473-3099(16)30127-X

14. Scherpenzeel CGM, Tijs SHW, den Uijl IEM, Santman-Berends IMGA, Velthuis AGJ, Lam TJGM. Farmers’ attitude toward the introduction of selective dry cow therapy. J Dairy Sci [Internet]. American Dairy Science Association; 2016 Oct 1 [cited 2017 Dec 18];99(10):8259–66. Available from: http://linkinghub.elsevier.com/retrieve/pii/S0022030216304623. https://doi.org/10.3168/jds.2016-11349

15. Cameron M, McKenna SL, MacDonald KA, Dohoo IR, Roy JP, Keefe GP. Evaluation of selective dry cow treatment following on-farm culture: risk of postcalving intramammary infection and clinical mastitis in the subsequent lactation. J Dairy Sci [Internet]. Elsevier; 2014 Jan;97(1):270–84. Available from: http://linkinghub.elsevier.com/retrieve/pii/S0022030213007352. https://doi.org/10.3168/jds.2013-7060

16. Council NR. Nutrient Requirements of Dairy Cattle [Internet]. Washington, D.C.: National Academies Press; 2001 [cited 2017 Apr 18]. Available from: http://www.nap.edu/catalog/9825.

17. Barnum DA, Newbould FH. The Use of the California Mastitis Test for the Detection Of Bovine Mastitis. Can Vet J = La Rev Vet Can [Internet]. 1961 Mar [cited 2017 Apr 19];2(3):83–90. Available from: https://www.ncbi.nlm.nih.gov/pmc/articles/PMC1585631/pdf/canvetj00173-0009.pdf. PMID: 17421323

18. Schalm OW, Noorlander DO. Experiments and observations leading to development of the California mastitis test. J Am Vet Med Assoc [Internet]. 1957 Mar 1 [cited 2017 Apr 19];130(5):199–204. Available from: http://www.ncbi.nlm.nih.gov/pubmed/13416088. PMID: 13416088

19. National Mastitis Council. Guidelines for evaluating teat skin condition. Natl Mastit Counc [Internet]. 2007 [cited 2017 Apr 19];1–5. Available from: http://www.nmconline.org/wp-content/uploads/2016/09/Guidelines-for-Evaluating.pdf.

20. Bentley Instruments. Somacount 2000 Operator’s Manual. Chaska; 1995. 12 p.

21. Cameron M, Keefe GP, Roy J-P, Stryhn H, Dohoo IR, McKenna SL. Evaluation of selective dry cow treatment following on-farm culture: Milk yield and somatic cell count in the subsequent lactation. J Dairy Sci [Internet]. American Dairy Science Association; 2015 Apr;98(4):2427–36. Available from: http://linkinghub.elsevier.com/retrieve/pii/S0022030215000570. https://doi.org/10.3168/jds.2014-8876

22. Philpot WN, Nickerson SC. Mastitis: Counter Attack [Internet]. Babson Bros. Company, editor. 1991 [cited 2017 Apr 19]. 150 p. Available from: https://books.google.co.in/books/about/Mastitis.html?id=OSHpGAAACAAJ.

23. Shook GE. Genetic Improvement of Mastitis Through Selection on Somatic Cell Count. Vet Clin North Am Food Anim Pract [Internet]. 1993;9(3):563–77. Available from: http://linkinghub.elsevier.com/retrieve/pii/S0749072015306228. https://doi.org/10.1016/S0749-0720(15)30622-8

24. Garcia ML, Moreno B, Bergdoll MS. Characterization of staphylococci isolated from mastitic cows in Spain. Appl Environ Microbiol [Internet]. 1980 Mar;39(3):548–53. Available from: http://www.pubmedcentral.nih.gov/articlerender.fcgi?artid=291376&tool=pmcentrez&rendertype=abstract. PMID: 7387155

25. Rodrigues E, Gonçalves AM, Silva W, Régis K, Móta TR. Perfil de sensibilidade antimicrobiana.2012;701–11.

26. Teixeira LM, Siqueira G, Shewmaker PL, Facklam RR. Manual of Clinical Microbiology [Internet]. 10th ed. Versalovic J, Jorgensen JH, Funke G, Warnock DW, Landry ML, Carroll KC, editors. Manual of Clinical Microbiology, 10th edition. American Society of Microbiology; 2011. 350-364 p. Available from: http://www.asmscience.org/content/book/10.1128/9781555816728. https://doi.org/10.1128/9781555816728

27. Lapage SP. Biochemical Tests for Identification of Medical Bacteria. In: Journal of Clinical Pathology [Internet]. Williams and Wilkins; 1976. p. 958–958. Available from: http://jcp.bmj.com/cgi/doi/10.1136/jcp.29.10.958-c. https://doi.org/10.1136/jcp.29.10.958-c

28. Peterkin PI. Compendium of methods for the microbiological examination of foods (3rd edn). Trends in Food Science & Technology. 1993. p. 199. https://doi.org/10.1016/0924-2244(93)90131-S

29. Equipe Estatcamp. Software Action. [Internet]. São Carlos - SP, Brasil: Estatcamp - Consultoria em estatística e qualidade; 2014. p. v3.2.60. Available from: www.portalaction.com.br.

30. Cameron M, Keefe GP, Roy JP, Dohoo IR, MacDonald KA, McKenna SL. Evaluation of a 3M Petrifilm on-farm culture system for the detection of intramammary infection at the end of lactation. Prev Vet Med [Internet]. Elsevier B.V.; 2013;111(1–2):1–9. https://doi.org/10.1016/j.prevetmed.2013.03.006

31. Bhutto AL, Murray RD, Woldehiwet Z. California mastitis test scores as indicators of subclinical intra-mammary infections at the end of lactation in dairy cows. Res Vet Sci [Internet]. Elsevier Ltd; 2012 Feb;92(1):13–7. Available from: http://dx.doi.org/10.1016/j.rvsc.2010.10.006. https://doi.org/10.1016/j.rvsc.2010.10.006

32. Mahmmod YS, Toft N, Katholm J, Grønbæk C, Klaas IC. Bayesian estimation of test characteristics of real-time PCR, bacteriological culture and California mastitis test for diagnosis of intramammary infections with Staphylococcus aureus in dairy cattle at routine milk recordings. Prev Vet Med [Internet]. Elsevier B.V.; 2013;112(3–4):309– Available from: http://dx.doi.org/10.1016/j.prevetmed.2013.07.021. https://doi.org/10.1016/j.prevetmed.2013.07.021

33. Bijl E, van Valenberg HJF, Huppertz T, van Hooijdonk ACM. Protein, casein, and micellar salts in milk: Current content and historical perspectives. J Dairy Sci [Internet]. Elsevier; 2013;96(9):5455–64. Available from: http://linkinghub.elsevier.com/retrieve/pii/S0022030213004931. https://doi.org/10.3168/jds.2012-6497

34. Gindonis V, Taponen S, Myllyniemi A-L, Pyörälä S, Nykäsenoja S, Salmenlinna S, et al. Occurrence and characterization of methicillin-resistant staphylococci from bovine mastitis milk samples in Finland. Acta Vet Scand [Internet]. 2013;55(1):61. https://doi.org/10.1186/1751-0147-55-61

35. Litwińczuk Z, Król J, Brodziak A. Factors determining the susceptibility of cows to 1. mastitis and losses incurred by producers due to the disease - A review. Ann Anim Sci [Internet]. 2015;15(4):819–31. Available from: http://www.degruyter.com/view/j/aoas.ahead-of-print/aoas-2015-0035/aoas-2015-0035.xml. https://doi.org/10.1515/aoas-2015-0035

36. Matyi SA, Dupre JM, Johnson WL, Hoyt PR, White DG, Brody T, et al. Isolation and characterization of Staphylococcus aureus strains from a Paso del Norte dairy1. J Dairy Sci [Internet]. 2013;96(6):3535–42. Available from: http://linkinghub.elsevier.com/retrieve/pii/S0022030213003032. https://doi.org/10.3168/jds.2013-6590

37. Blagitz MG, Souza FN, Batista CF, Diniz SA, Azevedo LFF, Silva MX, et al. Flow cytometric analysis: Interdependence of healthy and infected udder quarters. J Dairy Sci [Internet]. American Dairy Science Association; 2015;98(4):2401–8. Available from: http://linkinghub.elsevier.com/retrieve/pii/S0022030215000909. https://doi.org/10.3168/jds.2014-8727

38. Gonçalves JL, Tomazi T, Barreiro JR, Beuron DC, Arcari MA, Lee SHI, et al. Effects of bovine subclinical mastitis caused by Corynebacterium spp. on somatic cell count, milk yield and composition by comparing contralateral quarters. Vet J [Internet]. 2016 Aug;209:87–92. Available from: http://linkinghub.elsevier.com/retrieve/pii/S1090023315003330. https://doi.org/10.3168/jds.2014-8727

39. Condas LAZ, De Buck J, Nobrega DB, Carson DA, Roy J-P, Keefe GP, et al. Distribution of non-aureus staphylococci species in udder quarters with low and high somatic cell count, and clinical mastitis. J Dairy Sci [Internet]. American Dairy Science Association; 2017 Jul;100(7):5613–27. https://doi.org/10.3168/jds.2016-12479

40. Guarín JF, Ruegg PL. Short communication: Pre- and postmilking anatomical 1. characteristics of teats and their associations with risk of clinical mastitis in dairy cows. J Dairy Sci [Internet]. 2016 Oct [cited 2017 May 7];99(10):8323–9. Available from: http://linkinghub.elsevier.com/retrieve/pii/S0022030216304830. https://doi.org/10.3168/jds.2015-10093

41. Martins CMMR, Pinheiro ESC, Gentilini M, Benavides ML, Santos MV. Efficacy of a high free iodine barrier teat disinfectant for the prevention of naturally occurring new intramammary infections and clinical mastitis in dairy cows. J Dairy Sci [Internet]. American Dairy Science Association; 2017 Feb;100(5):3930–9. Available from: http://linkinghub.elsevier.com/retrieve/pii/S0022030217301571. https://doi.org/10.3168/jds.2016-11193

42. Vasquez AK, Nydam D V, Foditsch C, Wieland M, Lynch R, Eicker S, et al. Use of a culture-independent on-farm algorithm to guide the use of selective dry-cow antibiotic therapy. J Dairy Sci [Internet]. American Dairy Science Association; 2018 Jun;101(6):5345–61. Available from: http://linkinghub.elsevier.com/retrieve/pii/S0022030218302844. https://doi.org/10.3168/jds.2017-13807

43. Hadrich JC, Wolf CA, Lombard J, Dolak TM. Estimating milk yield and value losses from increased somatic cell count on US dairy farms. J Dairy Sci [Internet]. American Dairy Science Association; 2018;1–9. Available from: http://linkinghub.elsevier.com/retrieve/pii/S0022030218300389. https://doi.org/10.3168/jds.2017-13840

44. Vasquez AK, Nydam DV, Capel MB, Eicker S, Virkler PD. Clinical outcome comparison of immediate blanket treatment versus a delayed pathogen-based treatment protocol for clinical mastitis in a New York dairy herd. J Dairy Sci [Internet]. 2017 Apr [cited 2018 May 27];100(4):2992–3003. Available from: http://linkinghub.elsevier.com/retrieve/pii/S0022030217300905. https://doi.org/10.3168/jds.2017-13840

45. Scherpenzeel CGM, den Uijl IEM, van Schaik G, Riekerink RGMO, Hogeveen H, Lam TJGM. Effect of different scenarios for selective dry-cow therapy on udder health, antimicrobial usage, and economics. J Dairy Sci [Internet]. Elsevier; 2016 May 1 [cited 2017 Dec 18];99(5):3753–64. Available from: http://linkinghub.elsevier.com/retrieve/pii/S0022030216300078. https://doi.org/10.3168/jds.2015-9963

46. Sargeant JM, Leslie KE, Shirley JE, Pulkrabek BJ, Lim GH. Sensitivity and Specificity of Somatic Cell Count and California Mastitis Test for Identifying Intramammary Infection in Early Lactation. J Dairy Sci [Internet]. 2001 Sep [cited 2017 May 4];84(9):2018–24. Available from: http://linkinghub.elsevier.com/retrieve/pii/S0022030201746450. https://doi.org/10.3168/jds.S0022-0302(01)74645-0

47. Reardon S. Phage therapy gets revitalized. Nature [Internet]. 2014 Jun 5 [cited 2018 Jan 1];510(7503):15–6. Available from: https://amr-review.org/sites/default/files/AMR Review Paper - Tackling a crisis for the health and wealth of nations_1.pdf. https://doi.org/10.1038/510015a

48. Brogan DM, Mossialos E. A critical analysis of the review on antimicrobial resistance report and the infectious disease financing facility. Globalization and Health. 2016. https://doi.org/10.1186/s12992-016-0147-y

49. Fitchett JR, Atun R. Antimicrobial resistance: Opportunity for Europe to establish global leadership. Lancet Infect Dis [Internet]. 2016 Apr [cited 2018 Jan 1];16(4):388–9. Available from: http://linkinghub.elsevier.com/retrieve/pii/S1473309915004107. https://doi.org/10.1016/S1473-3099(15)00410-7

50. Bragginton EC, Piddock LJV. UK and European Union public and charitable funding from 2008 to 2013 for bacteriology and antibiotic research in the UK: An observational study. Lancet Infect Dis [Internet]. Elsevier; 2014 Sep 1 [cited 2018 Jan 1];14(9):857–68. Available from: http://www.ncbi.nlm.nih.gov/pubmed/25065509. https://doi.org/10.1016/S1473-3099(14)70825-4

